# Potential transmission chains of variant B.1.1.7 and co-mutations of SARS-CoV-2

**DOI:** 10.1101/2021.04.16.440141

**Authors:** Jingsong Zhang, Yang Zhang, Junyan Kang, Shuiye Chen, Yongqun He, Benhao Han, Mofang Liu, Lina Lu, Li Li, Zhigang Yi, Luonan Chen

**Author notes:** Correspondence: Luonan Chen or Zhigang Yi. These authors contributed equally: Jingsong Zhang, Yang Zhang.

## Abstract

The presence of SARS-CoV-2 mutants, including the emerging variant B.1.1.7, has raised great concerns in terms of pathogenesis, transmission, and immune escape. Characterizing SARS-CoV-2 mutations, evolution, and effects on infectivity and pathogenicity is crucial to the design of antibody therapies and surveillance strategies. Here we analyzed 454,443 SARS-CoV-2 spike genes/proteins and 14,427 whole-genome sequences. We demonstrated that the early variant B.1.1.7 may not have evolved spontaneously in the United Kingdom or within human populations. Our extensive analyses suggested that Canidae, Mustelidae or Felidae, especially the Canidae family (for example, dog) could be a possible host of the direct progenitor of variant B.1.1.7. An alternative hypothesis is that the variant was simply yet to be sampled. Notably, the SARS-CoV-2 whole genome represents a large number of potential co-mutations with very strong statistical significances (p value<E– 44). In addition, we used an experimental SARS-CoV-2 reporter replicon system to introduce the dominant co-mutations NSP12_c14408t, 5’UTR_c241t, and NSP3_c3037t into the viral genome, and to monitor the effect of the mutations on viral replication. Our experimental results demonstrated that the co-mutations significantly attenuated the viral replication. The study provides valuable clues for discovering the transmission chains of variant B.1.1.7 and understanding the evolutionary process of SARS-CoV-2.

## Introduction

Since the outbreak in December 2019, COVID-19 has been pandemic in over 200 countries. Cases of infection and mortalities have been surging and are an ongoing threat to public health^1,2^. COVID-19 is caused by infection with the novel coronavirus SARS-CoV-2^3–5^. Although as a coronavirus, SARS-CoV-2 has genetic proofreading mechanisms^6–8^, the persistent natural selection pressure in the population drives the virus to gradually accumulate favorable mutations^6,9,10^. Much attention has been paid to the mutations and evolution of SARS-CoV-2^11–15^, since mutations are related to the infectivity and pathogenicity of viruses^16–21^. Beneficial mutants of the virus can better evolve and adapt to the host^9^, either strengthening or weakening the infectivity and pathogenicity. In addition, certain variants may generate drug resistance and reduce the efficacy of vaccines and therapeutics^22–26^. In short, studying mutations and evolution in detail is vital to understand the transformations of viral properties and to control the pandemic.

A new variant of SARS-CoV-2 named VOC-202012/01 (Variant of Concern 202012/01) or lineage B.1.1.7 was first detected in the United Kingdom last December^27^. It appears to be substantially more transmissible than other variants^28^. The variant has been growing exponentially in the United Kingdom and rapidly spreading to other countries^29,30^. However, it is not yet clear if it evolved spontaneously in the United Kingdom or was imported from other countries. Studying how the variant B.1.1.7 mutates can enable researchers to track its spread over time and to understand the evolution of SARS-CoV-2.

In this study, large-scale SARS-CoV-2 sequences, consisting of more than 454,000 spike genes/proteins and 14,000 whole-genome sequences were analyzed. Our extensive sequence analysis showed that many mutations always co-occur not only in the spike protein of B.1.1.7, but in the whole genome of SARS-CoV-2. The mutation trajectories of the spike protein indicate that the early variant B.1.1.7 did not evolve spontaneously in the United Kingdom or even within human populations. We also investigated possible SARS-CoV-2 transmission chains of the variant B.1.1.7 based on the mutation analysis of large-scale spike proteins and the cluster analysis of spike genes. Over the whole genome, the top 25 high-frequency mutations of SARS-CoV-2 converged into several potential co-mutation patterns, each of which showed a strong correlation with a very strong statistical significance (p value<E–44). The potential co-mutations depicted the evolutionary trajectory of SARS-CoV-2 virus in the population, shaping variable replication of SARS-CoV-2. In addition, we further explored the effect of the dominant (co-)mutations 5’UTR_c241t, NSP3_c3037t, and NSP12_c14408t on viral replication using a SARS-CoV-2 replicon based on a four plasmid *in-vitro* ligation system. The results suggest that such mutations significantly attenuate the replication of SARS-CoV-2.

## Results

### Evolutionary trajectories of variant B.1.1.7

The variant B.1.1.7 was generally defined by multiple amino acid changes including 3 deletions (69-70del and 145del) and 7 mutations (N501Y, A570D, D614G, P681H, T716I, S982A, and D1118H) in the spike protein^31^. The number of non-adjacent co-occurrent changes indicates that they resulted from accumulated mutations. We therefore explored the evolutionary trajectories of B.1.1.7 by tracing the incremental mutations (Fig. 1a). All routes along the directions of the arrows are possible evolutionary trajectories of lineage B.1.1.7. Among all the mutation routes, the green one was the most probable mutation trajectory based on the number of variant strains. However, it was unlikely that the earliest variant B.1.1.7 (GISAID: EPI_ISL_601443, 2020-09-20, England) with 9 mutations evolved from the existing variants with 3–8 mutations, because the former arose much earlier than the latter. More than 454,000 SARS-CoV-2 strains have been collected and extensive sequenced from infected humans without finding intermediate variants with 3–9 mutations. It is therefore unlikely that the intermediate variants with 3–8 mutations have infected humans. Thus, the early variant B.1.1.7 might not have arisen spontaneously in the UK or within human populations. An alternative hypothesis is that spillover likely occurred from susceptible animals.

**Fig. 1.**
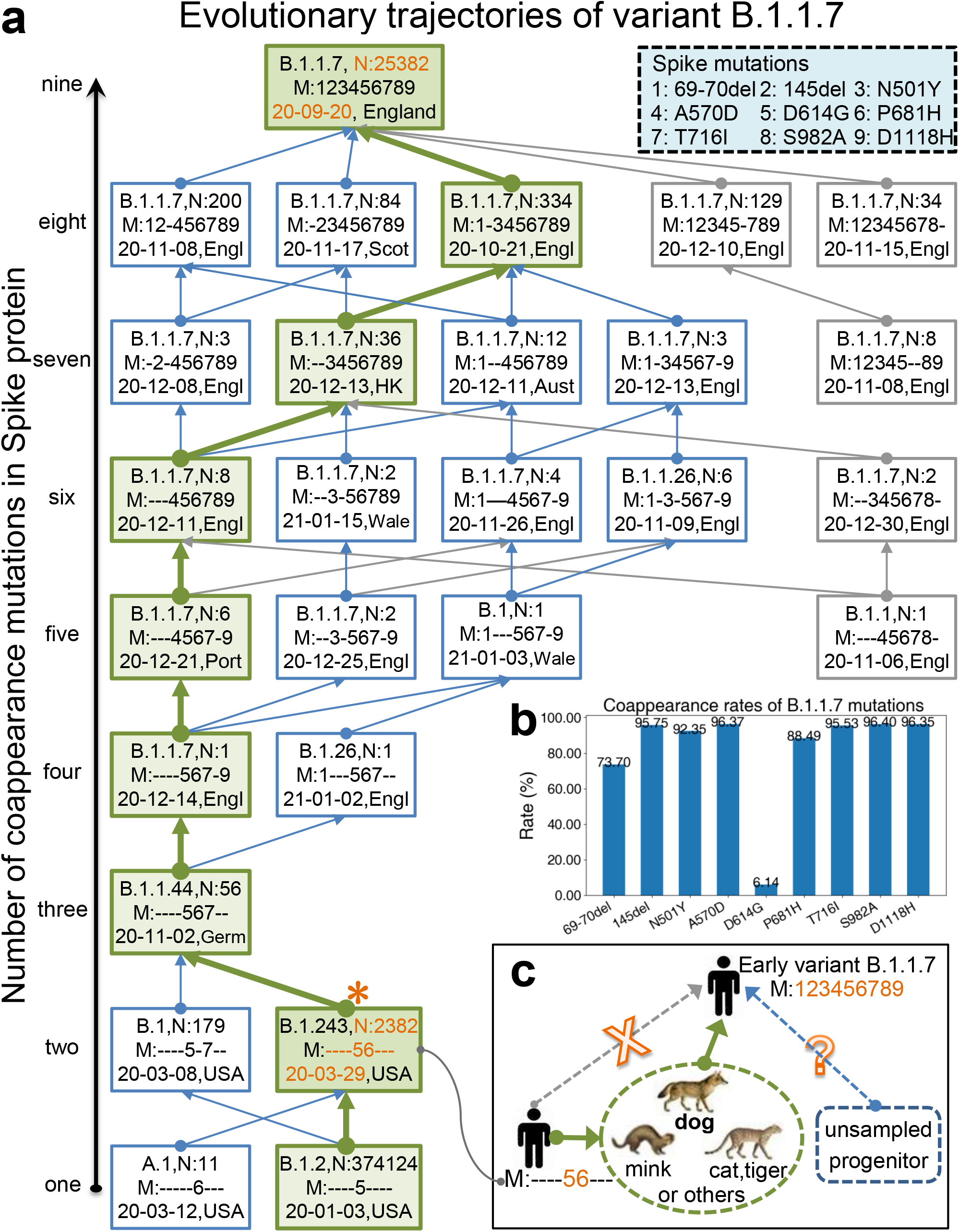
Evolutionary trajectories of variant B.1.1.7. **a** Incremental mutations of variant B.1.1.7. The digits in the upper-right-corner rectangle with dotted line indicate the labels of mutations. For simplicity, the 69–70 deletions were labeled as “1”, and the other mutations “2”-“9” respectively. The bottom nodes (rectangles) represent the variants with one mutation and the top one was the early variant B.1.1.7. Each rectangle with solid line consists of lineage (e.g., B.1.243), number of strains (e.g., N:2382), mutation sites (e.g., M:----56---), the earliest collection date (e.g., 20-03-29, i.e., 2020-03-29), and collection location (e.g., USA). In the labels of the mutation sites, sign “-” indicated the corresponding site did not mutate. All routes along the directions of the arrows are possible evolutionary trajectories of lineage B.1.1.7, where the green one was the most probable mutation trajectory. Large-scale SARS-CoV-2 analysis demonstrates that the early variant B.1.1.7 might not have arisen spontaneously in the UK or within human hosts. **b** Coappearances of variant B.1.1.7 mutations. At least five mutations form a potential co-mutation pattern (coappearance rate > 95%). **c** Possible transmission chains of variant B.1.1.7. Canidae, Mustelidae or Felidae, especially the Canidae family (for example, dog) could be a possible host of the direct progenitor of variant B.1.1.7.

The co-appearance rates (see Materials and Methods) of all nine mutations are shown in Fig. 1b. We found that at least five mutations (145del, A570D, T716I, S982A, and D1118H) of variant B.1.1.7 significantly co-occurred (rate>95%), which indicates a potential co-mutation pattern in the spike protein, causing us to wonder what selection pressure drove such co-occurrences of mutations and rapid evolution in the population of SARS-CoV-2. Note that coronaviruses generally tend to exhibit rapid evolution when they jump to a different species^32^. We therefore analyzed the key spike genes and proteins of existing SARS-CoV-2 strains collected from animals to find a possible direct progenitor of variant B.1.1.7. The variant with mutations “56” (labeled by “*” in Fig. 1a, termed star variant) had the minimum phylogenetic distance with EPI_ISL_699508, which was collected from a dog on 2020-07-28 (Fig. 2) using MEGA^33,34^ (see Materials and Methods). The strains collected from tigers, minks, and cats were also close to the star variant. Our extensive analyses including mutations, phylogeny (Fig. 2), collection date/location and the number of sequences (Tables S1-S3) suggested that Canidae, Mustelidae or Felidae, especially the Canidae family (for example, dog) could be a possible host of the direct progenitor of variant B.1.1.7. The possible transmission chains of variant B.1.1.7 are shown in Fig. 1c. This star variant strains in humans could not have evolved into the early variant B.1.1.7, but they might have infected high-density yet susceptible animals (such as dogs) and adapted to these species through rapid mutation. Such progenitor variants comprised most or all of the mutations of the early variant B.1.1.7 within the Canidae family populations, and they may have spilled back to humans after the rapid mutation period.

**Fig. 2.**
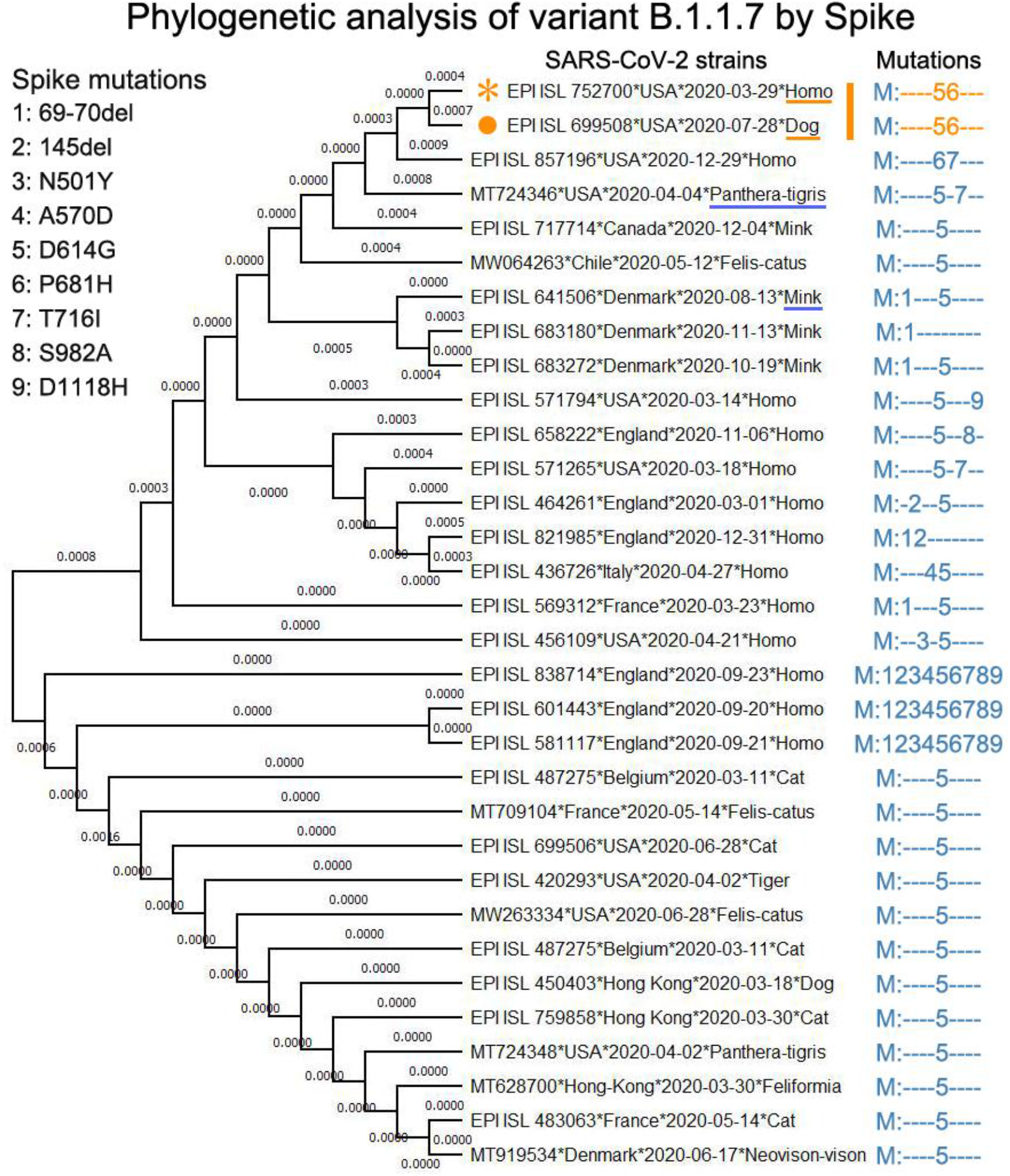
The Canidae family could be a possible host of the direct progenitor of variant B.1.1.7. The digits on the left of the figure indicate the labels of mutations, which correspond with the mutation labels in Fig. 1a. The strains shown in the center of the figure contain at least one spike mutation of variant B.1.1.7. And these strain examples cover all existing SARS-CoV-2 viruses that collected from animal hosts. The strain labeled by orange star corresponds with the star variant in Fig. 1a. The strain with orange solid-round label was collected from a dog on 2020-07-28. Such two strains share the same mutations “56” and have the minimum phylogenetic distance by MEGA tool. Canidae, Mustelidae or Felidae, especially the Canidae family (for example, dog) could be a possible host of the direct progenitor of variant B.1.1.7 based on existing stains collected before the end of Jan. 2021.

### High-frequency mutations converge into potential co-mutations

Based on sequence alignment and mutation analysis, we found that 7,441 nucleotide alterations in the viral 29903-letter RNA code occurred at least once in the samples from COVID-19 patients. These mutations were dispersed in the 14,427 SARS-CoV-2 strains collected from all around the world. As shown in the heatmap of the top 1% high-frequency mutations (Table S4), some sites show very similar mutation rates on most days in samples isolated globally (Fig. S1), including 8,898 and 815 samples isolated from the U.S. (Fig. S2) and Australia (Fig. S3). Therefore, these mutations shown in Fig. S4a were selected and clustered into co-occurrences, which we called potential co-mutation patterns. From the landscape of the mutation rates (Fig. S4a), 25 nucleotide sites were clearly clustered into several potential co-mutation patterns. Among these patterns, there was one consisting of the top 4 high-frequency mutations (i.e., 5’UTR_c241t, NSP3_c3037t, NSP12_c14408t, and S_a23403g), which converged into a dominant potential co-mutation pattern. Such co-occurrence lineage has been found in almost all sequenced samples of SARS-CoV-2. Within this co-occurrence pattern, mutation S_ a23403g resulted in the amino acid change (D614G) that apparently enhances viral infectivity^6,35^, albeit debate exists^16^. Notably, there were three successive sites at the 28881^st^ to 28883^rd^ positions of the virus (N_g28881a, N_g28882a, and N_g28883c) that strictly co-occurred. Comparing Fig. S4a-c and Table S4, we found that the top 14 high-frequency mutations formed five common co-occurrence patterns.

To assess the above co-occurrence patterns, we analyzed the correlations and statistical significance levels of the high-frequency co-occurrence mutations. The heatmap of the paired Pearson-correlation-coefficients (Fig. 3a) shows that the top 25 high-frequency mutations clearly cluster into several potential co-mutation groups/patterns with very strong correlation (≥0.8). By regression analyses, the above co-occurrence patterns have statistical significance levels with p values less than 10^−44^ (Fig. 3b). The detailed mutation transitions (Fig. 3c–k, Figs. S5–7) provide further evidence that the above mutations form co-mutation patterns.

**Fig. 3.**
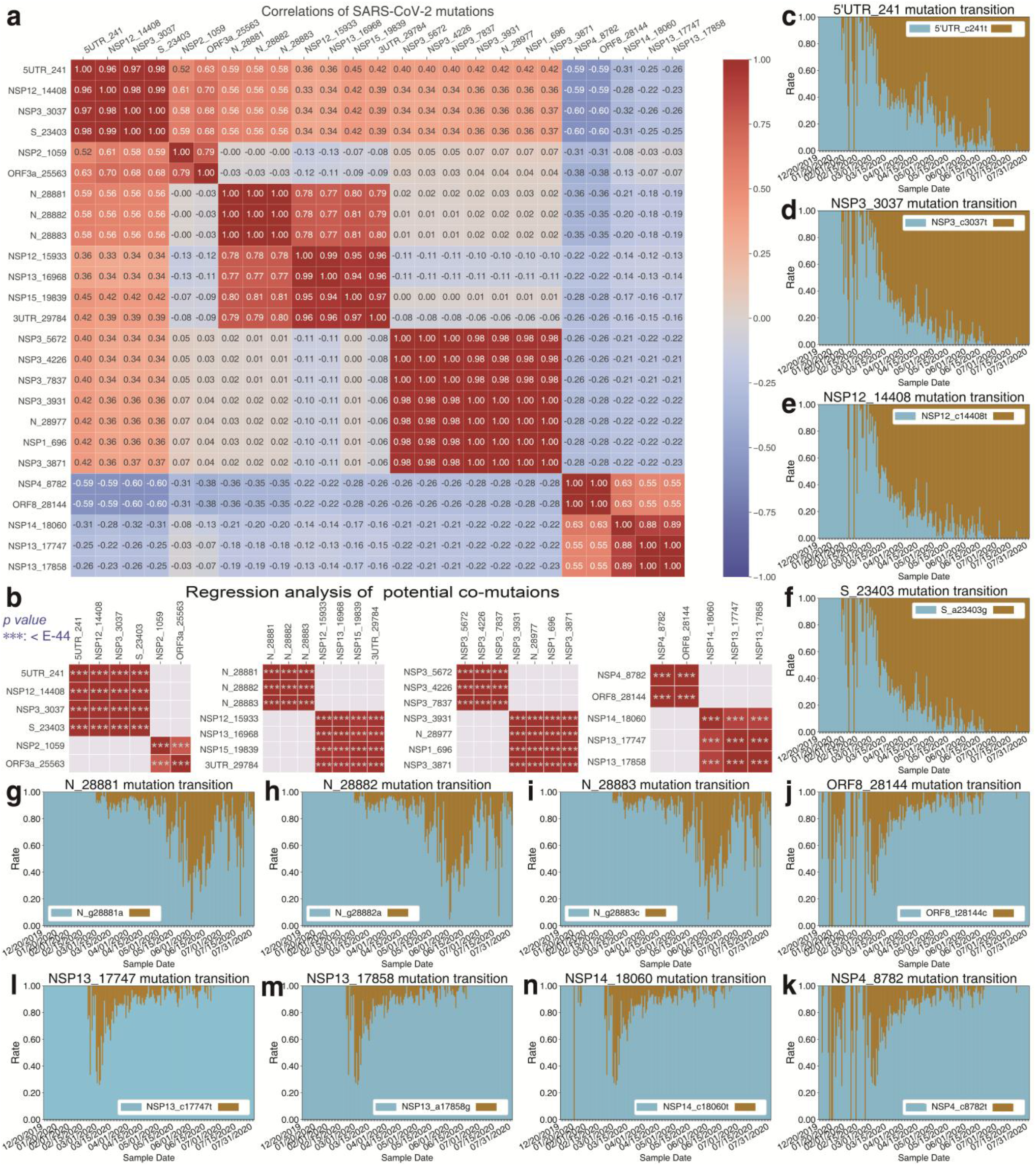
The strong correlations suggest that the top 25 mutations form eight potential co-mutation patterns. **a** The correlation heatmap of the top 25 mutations. These mutations could be grouped into several clusters with high Pearson-correlation-coefficient (PCC). **b** Regression analysis of mutations shows that eight clusters all denote the statistical significance level: ***p value < E-44. c to k show the transitions of the high-frequency mutations. The sky-blue represents the rate per day of initial residue in population and the golden the rate per day of substitution/mutant. These mutation transitions provide further evidence that the above mutations potentially form co-mutation patterns.

### Dominant mutations attenuate viral replication

We further explored the effect of the dominant mutations 5’UTR_c241t, NSP3_c3037t, and NSP12_c14408t on viral replication using a SARS-CoV-2 replicon based on a four-plasmid *in-vitro* ligation system. This replicon is devoid of the viral structural proteins while undergoing viral replication, and the viral replication is sensitive to the antiviral agent remdesivir^36^. The 5’UTR_c241t mutation resides in a highly conserved region in the 5’UTR (Fig. 4a). The NSP3_c3037t mutation is synonymous. The NSP12_c14408t mutation is nonsynonymous with an amino acid change of a conserved amino acid P323 in the viral RNA-dependent RNA polymerase (Fig. 4b). We introduced the NSP12_c14408t mutation or the NSP12_c14408t mutation with the other two mutations 5’UTR_c241t and NSP3_c3037t into the replicon plasmids. The fragments were released from the plasmids by BsaI digestion, and then assembled by *in-vitro* ligation with T4 ligase (Fig. 4c). Replicon RNA transcribed from the ligation products was co-transfected with N mRNA into Huh7 cells. RNA replication was monitored by measuring the secreted *Gaussia* luciferase activity in the supernatants. Enzymatic dead mutants (759-SAA-761) of the RNA-dependent RNA polymerase NSP12 were introduced, and the mutated replicon served as a non-replication control. As shown in Fig. 4d, transfection of WT replicon RNA resulted in an obvious increase of luciferase activity, and SAA RNA did not replicate as expected. Introduction of NSP12_c14408t mutation resulted in a significant reduction of viral replication. The combination of NSP12_c14408t mutation with the other two mutations further significantly but only marginally reduced viral replication. These results demonstrate that the P323L mutation in the viral RNA-dependent RNA polymerase reduces viral replication, and the synonymous mutations may further attenuate viral replication.

**Fig. 4.**
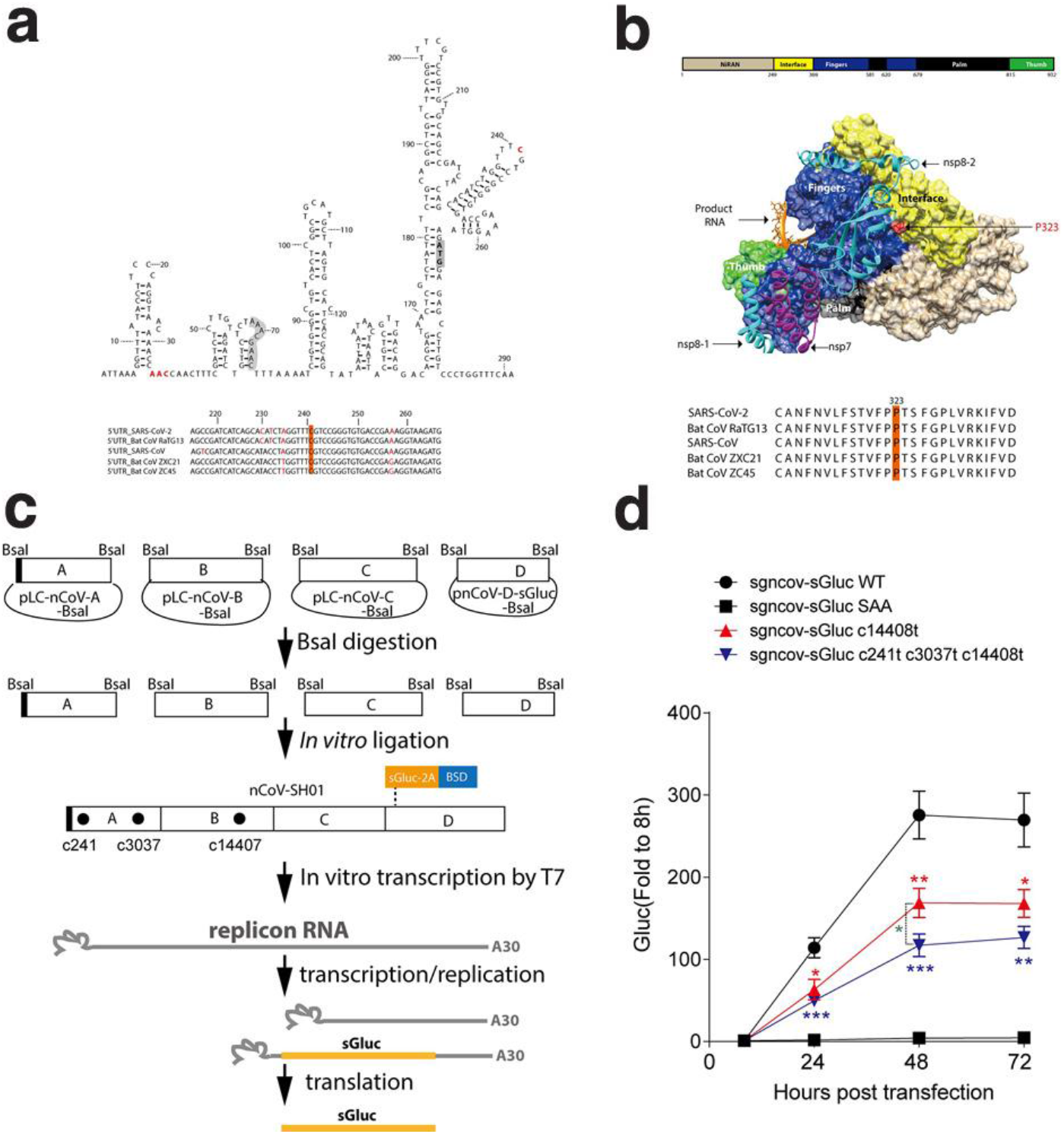
Dominant co-mutation attenuates viral replication. **a** Predicted RNA structure of the SARS-CoV-2 5’UTR. RNA structure of the 400-nt 5’UTR was predicted by “RNAstructure” (http://rna.urmc.rochester.edu/RNAstructureWeb). The start codon for nsp1 is grey, the TRS-L is orange, and the mutated nucleotides are red. The bottom panel shows the alignment of the 5’UTR of SARS-CoV-2 with 5’UTRs of related viruses, with c241 highlighted. **b** Structure of SARS-CoV-2 RdRp/RNA complex. The structure of SARS-CoV-2 RdRp/RNA complex (PDB, 6X2G) was visualized by Chimera (UCSF). The P323 mutation is highlighted in red, with the alignment of the amino acid sequences of SARS-CoV-2 and related viruses near the P323 position. **c** Schematic of the *in-vitro* ligation system for SARS-CoV-2 replicon. Four plasmids encompassing the viral genome were digested by BsaI to release the four fragments. After gel purification, the fragments were ligated by T4 ligase. The ligation products were purified and used as template for RNA *in-vitro* transcription. sGluc, secreted *Gaussia* luciferase; 2A, foot-and-mouth disease virus (FMDV) 2A peptide; BSD, blasticidin. **d** Huh7 cells were co-transfected with *in-vitro* transcribed replicon RNA (WT or the indicated mutants) and an mRNA encoding the SARS-CoV-2 N protein. The luciferase activity in the supernatants was measured at the time points indicated. Medium was changed at 8 hours post-transfection. Data are shown as mean±SEM (n=8). SAA, the NSP12 polymerase active-site mutant. Unpaired Student’s t-test was performed between the mutants and wild type (WT) and between the mutants as indicated (statistical significance level: *p value<0.05, **p value<0.01, ***p value<0.001).

## Discussion

A well-resolved phylogeny of variant B.1.1.7 spike genes provides an opportunity to understand the evolutionary process and transmission chains of variant B.1.1.7. Our incremental mutation and phylogenetic analyses on large-scale SARS-CoV-2 spike proteins/genes revealed that the early variant B.1.1.7 might not have evolved spontaneously in the United Kingdom or within human populations. In this case the spillover likely occurred from susceptible animals. Current evidence^37–39^ indicates that SARS-CoV-2 can effectively infect both domestic animals (for example, dog, cat, pig and bovine) and wild animals (for example, mink, rabbit and fox) by binding their angiotensin converting enzyme 2 (ACE2). Our further analyses including mutations, phylogeny, collection date/location and the number of sequences suggested that the earliest variant B.1.1.7 possibly originated from Canidae, Mustelidae or Felidae, especially the Canidae family (for example, dog). The cases^40^ that the variant B.1.1.7 can easily infect dogs and cats indicated that both are susceptible to B.1.1.7. Still, due to the limited information available to date, an alternative hypothesis is that the direct progenitor of variant B.1.1.7 is yet to be sampled. In addition to variant B.1.1.7, as a future topic we will work on the analysis of other lineages such as P.1, B.1.351, B.1.427, and B.1.42, when sufficient numbers of their sequences are available.

By tracing the mutation trajectories, we found that at least five mutations of the spike proteins always co-occurred, and a large number of potential co-mutations appeared in the top 1% high-frequency mutations of SARS-CoV-2 whole genome. It has been documented that the mutation S_ a23403g results in the amino acid change of the spike protein D614G and enhances viral infectivity^19,41–44^. Here, by using a SARS-CoV-2 reporter replicon system, we demonstrated that the one of the dominant co-mutations NSP12_c14408t significantly reduced viral replication and combination of NSP12_c14408t mutation with the other two synonymous mutations 5’UTR_c241t and NSP3_c3037t although significantly but only marginally reduced viral replication further. As the 5’UTR play an important role in regulating viral replication, the synonymous mutations 5’UTR_c241t may attenuate viral replication by change RNA secondary structure^45^. These findings imply that SARS-CoV-2 undergoes an evolution toward enhancing viral infectivity while attenuating viral replication. SARS-CoV-2 has exhibited significant mutations and co-mutations. We evaluated the replication of a co-mutation pattern including three dominant mutations. If other mutations act similarly on the viral replication needs to be verified. These results can be further explored for efficient vaccine design in our future work. In summary, this study provides insights into the transmission chains of variant B.1.1.7 and the effect of viral dominant mutations on viral evolution.

## Materials and Methods

### Data selection and pre-processing

The 454,443 spike gene/protein sequences of SARS-CoV-2 were obtained at https://www.gisaid.org/. The NCBI website at https://www.ncbi.nlm.nih.gov/sars-cov-2/ has released more than 1.7 thousand sequences of SARS-CoV-2 viruses before July 31, 2020. We selected 14,427 sequences that satisfied two criteria: (1) having specific collection dates; (2) sequence-lengths being no less than 29,305 nt (29903*0.98). It is inevitable that some sites of sequences are equivocal owing to the limitation of sequencing depth. For instance, many sites were labeled as letter N in genome sequences. The noise of indeterminate nucleic-acids was taken into consideration in our experiments so as to boost accuracy. The co-mutation rate of multi-site co-mutations was calculated by 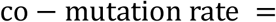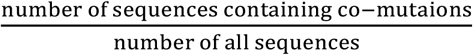. Moreover, the co-appearance rate of a mutation in B.1.1.7 variant was defined by 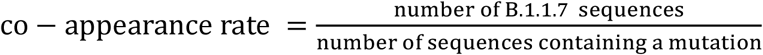.

### Possible animal host analyses

In addition to the phylogenetic analysis, we further explored the possible animal hosts of the direct progenitor of variant B.1.1.7 by mutations, collection time/space of strains, the number of sequences and the edit distance^46,47^ of mutations (Table S1-2). Due the late lockdown policies of some governmental agencies, the spread of SARS-CoV-2 has not been prevented well in Europe, America, and Australia. We could ignore the impact of policies for studying the origin of variant B.1.1.7. We quantified the multiple impact factors of viral transmission as shown in Table S3 based on the criterion that the smaller the value, the more similar. The results still supported that the Canidae family is a possible host of the direct progenitor of variant B.1.1.7.

### MEGA version and parameter settings

Version: MEGA-X

Statistical Method: Maximum Likelihood

Test of Phylogeny: None

Model/Method: Jones-Taylor-Thornton (JTT) model

Rates among Sites: Uniform Rates

Gaps/Missing Data Treatment: Use all sites

ML Heuristic Method: Nearest-Neighbor-Interchange (NNT)

Initial Tree for ML: Make initial tree automatically (Default - NJ/BioNJ)

Branch Swap Filter: None

Number of Threads: 7

### Statistical analysis

The Pearson-correlation-coefficient (PCC) is a classic statistic that measures linear correlation between two variables. Its value ranges from −1.0 to 1.0. Normally, the two variables meet a strong correlation or a very strong correlation when the absolutes value of PCC is between 0.6 and 0.8 or between 0.8 and 1.0. Linear regression is a linear approach to model the relationship between a scalar response and one or more variables. We used PCC and significance level (p value) of regression analysis to evaluate the relationships of the co-occurrence mutations in large-scale SARS-CoV-2 examples.

### Plasmids

Four plasmids encompassing the viral genome (pLC-nCoV-A-BsaI, pLC-nCoV-B-BsaI, pLC-nCoV-C-BsaI, and pnCoV-D-sGluc-BsaI) were described previously^36^. The 5’UTR_c-241-t and NSP3_c-3037-t mutations were introduced into the pLC-nCoV-A-BsaI by fusion PCR. The NSP12_c-14408-t mutation was introduced into the pLC-nCoV-B-BsaI by fusion PCR.

### Cell lines

The human hepatoma cells Huh 7 were purchased from the Cell Bank of the Chinese Academy of Sciences (www.cellbank.org.cn) and routinely maintained in Dulbecco’s modified medium supplemented with 10% FBS (Gibco) and 25 mM HEPES (Gibco).

### *In-vitro* ligation

BsaI digested fragments were gel purified using Gel Extraction Kit (OMEGA) and ligated with T4 ligase (New England Biolabs) at room temperature for 1 h. The ligation products were phenol/chloroform extracted, precipitated by absolute ethanol, and resuspended in nuclease-free water, quantified by determining the A260 absorbance.

### *In-vitro* transcription

Purified *in-vitro* ligated product was used as template for the *in-vitro* transcription by mMESSAGE mMACHINE T7 Transcription Kit (Ambion) according to the manufacturer’s protocol. For N mRNA production, we amplified the N coding region by PCR (sense: *GGC ACA CCC CTT TGG CTC T*; antisense: *TTT TTT TTT TTT TTT TTT TTT TTT TTT TTT TTT TTT TTT TCT AGG CCT GAG TTG AGT CAG CAC*) with phCMV-N as template. Then the purified PCR product was used as a template for *in-vitro* transcription by mMESSAGE mMACHINE T7 Transcription Kit as described above. RNA was purified by RNeasy mini Elute (Qiagen), eluted in nuclease-free water, and quantified by UV absorbance (260 nm).

### Transfection

Cells were seeded onto 48-well plates at a density of 7.5×10^4^ per well and then transfected with 0.3 μg *in-vitro* transcribed RNA using a TransIT-mRNA transfection kit (Mirus) according to the manufacturer’s protocol.

### Luciferase activity

Supernatants were taken from cell medium and mixed with equal volumes of 2×lysis buffer (Promega). Luciferase activity was measured with Renilla luciferase substrate (Promega) according to the manufacturer’s protocol.

## Acknowledgments

We thank associate prof. T.Z. and Dr. S.T.H. for useful comments on the manuscript. This work is supported by the National Key Research and Development Program of China (2017YFA0505500 to L.N.C., 2017YFC0909502 to J.S.Z.); the Strategic Priority Research Program of the Chinese Academy of Sciences (XDB38040400 to L.N.C.); National Science Foundation of China (31771476 and 31930022 to L.N.C, 61602460 and 11701379 to J.S.Z.); Shanghai Municipal Science and Technology Major Project (2017SHZDZX01 to L.N.C.); National Science and Technology Major Project of China (2017ZX10103009 to Z.G.Y.); Emergency Project of Shanghai Science and Technology Committee (20411950103 to Z.G.Y.); National Postdoctoral Program for Innovative Talent (BX20180331 to J.Y.K.); and China Postdoctoral Science Foundation (2018M642018 to J.Y.K.).

## Author contributions

L.N.C. and J.S.Z. designed the study. Z.G.Y. and J.S.Z. designed the experiments. J.S.Z. analyzed data. Y. Z. performed the experiments of viral replication. J.S.Z., Z.G.Y., and J.Y.K. designed the figures. S.C. repeated and checked the experiments of viral replication. H.B.H. checked the computational analyses. J.S.Z. and Z.G.Y. wrote the manuscript. Y.Q.H, M.F.L., L.N.L, and L.L. polished the manuscript. All authors participated in result interpretation and discussion.

## Data availability

The raw sequence data reported in this paper have been deposited in the GISAID and NCBI websites at https://www.gisaid.org/ and https://www.ncbi.nlm.nih.gov/sars-cov-2/, respectively. Code is available from the corresponding author on reasonable request.

