## Supplementary material for "Potential transmission chains of variant B.1.1.7 and co-mutations of SARS-CoV-2": SI

**This PDF file includes:**

Supplementary Text

Supplementary Figures. S1-7

Supplementary Table S1-7

### Supplementary Text : Evolutionary trajectories of potential co-mutations

We also studied the evolutionary trajectories of the potential co-mutations over time in detail. Comparing Fig. S4a with the main text Fig. 3a, the 25 nucleotide sites with high mutation rates could cluster into eight potential co-mutation patterns. The frequencies of mutations and potential co-mutations were shown in Fig. S6a-h. As expected, the top 4 high-frequency mutations converged into a dominant potential co-mutation pattern (green curve of Fig. S6a, main text Fig. 3c-f), and such pattern's lineage represented nearly 100% of the samples since early July, 2020. Notably, the potential co-mutations almost always rose in frequency since their appearances. We could also infer that such pattern's lineage will continually dominate all the samples from COVID-19 patients. Both ORF3a_g25563t and NSP2_c1059t were very high-frequency mutations (both more than 30%,Table S4) and the trajectory of the co-occurrences almost overlapped with that of NSP2_c1059t (Fig. S6b). This pattern maintained a medium co-occurrence rate on the whole. Fig. S6c showed the potential co-mutation pattern of three successive sites (N_28881, N_28882, and N_28883). Unlike the patterns shown in Fig. S6a and S6b, the co-occurrence rates of the three successive mutations (Fig. S6c) increased first and then decreased slightly. The trajectories of co-occurrence mutations in Fig. S6e and S6f shared a very similar trend that increased rapidly first, decreased next, and almost disappeared finally. The co-occurrence rates of Fig. S6d first rose, and then gradually dropped to near zero. Interestingly, the potential co-mutation patterns of Fig. S6g and S6h occurred when Fig. S6d's patterns disappeared. Fig. 6i illustrates the residue positions of the above eight potential co-mutation patterns. For these patterns, several patterns occurred within one gene (Fig. S6i(c), (h)) or between two adjacent genes (Fig. S6i(f)), but most of them spanned distant genes (Fig. S6i(a), (b), (d), (g), (e)). The long-range potential co-mutations may imply the folding of RNA spatial structure and the possible interactions between/among these sites.

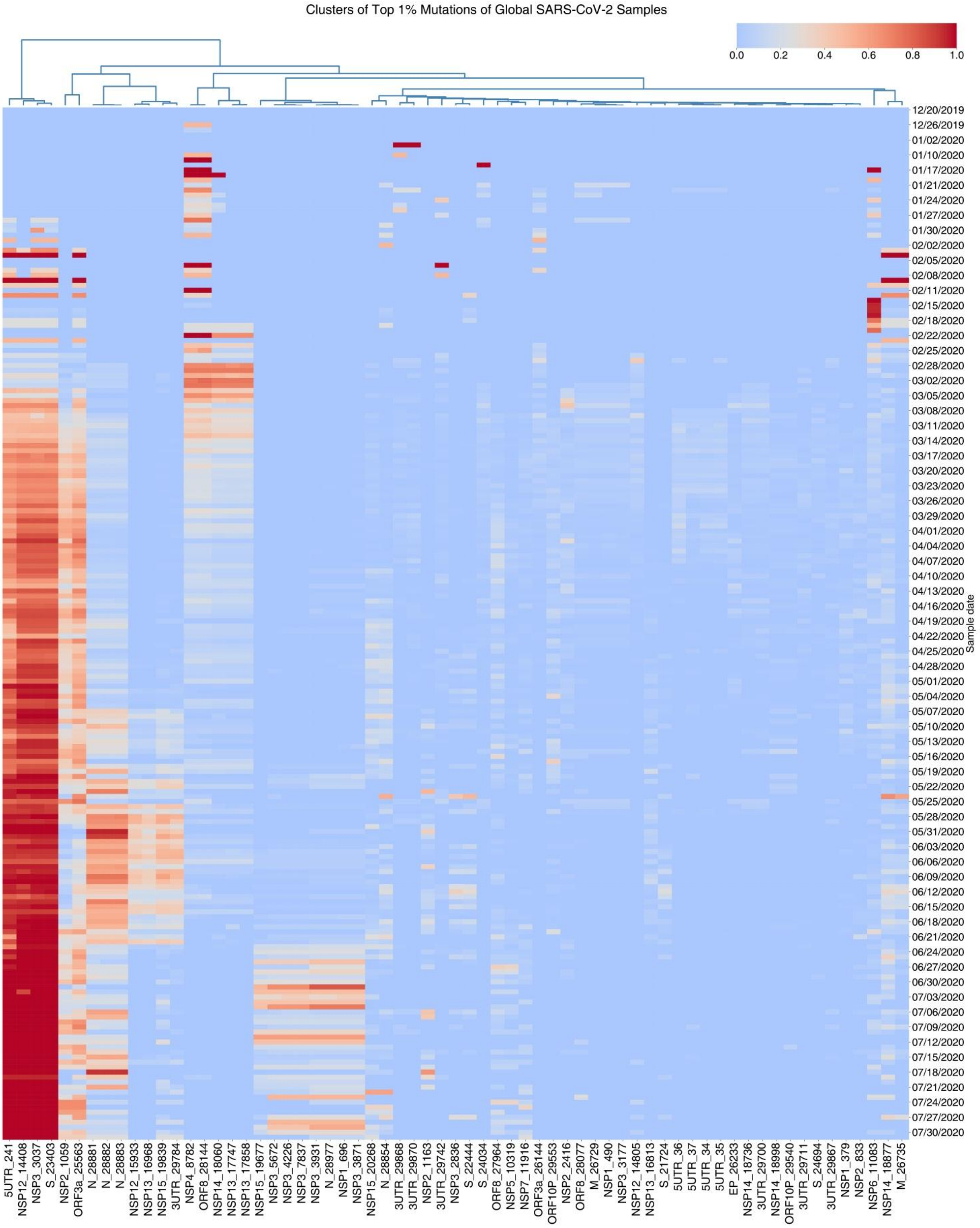


**Fig. S1 Ongoing mutations indicate potential co-mutations of SARS-CoV-2 from global samples.** The number of sequences per collection date is shown in Table S5. Top 1% high-frequency mutations consist of 65 sites. There are about 25 mutations clustered into several potential co-mutation patterns.

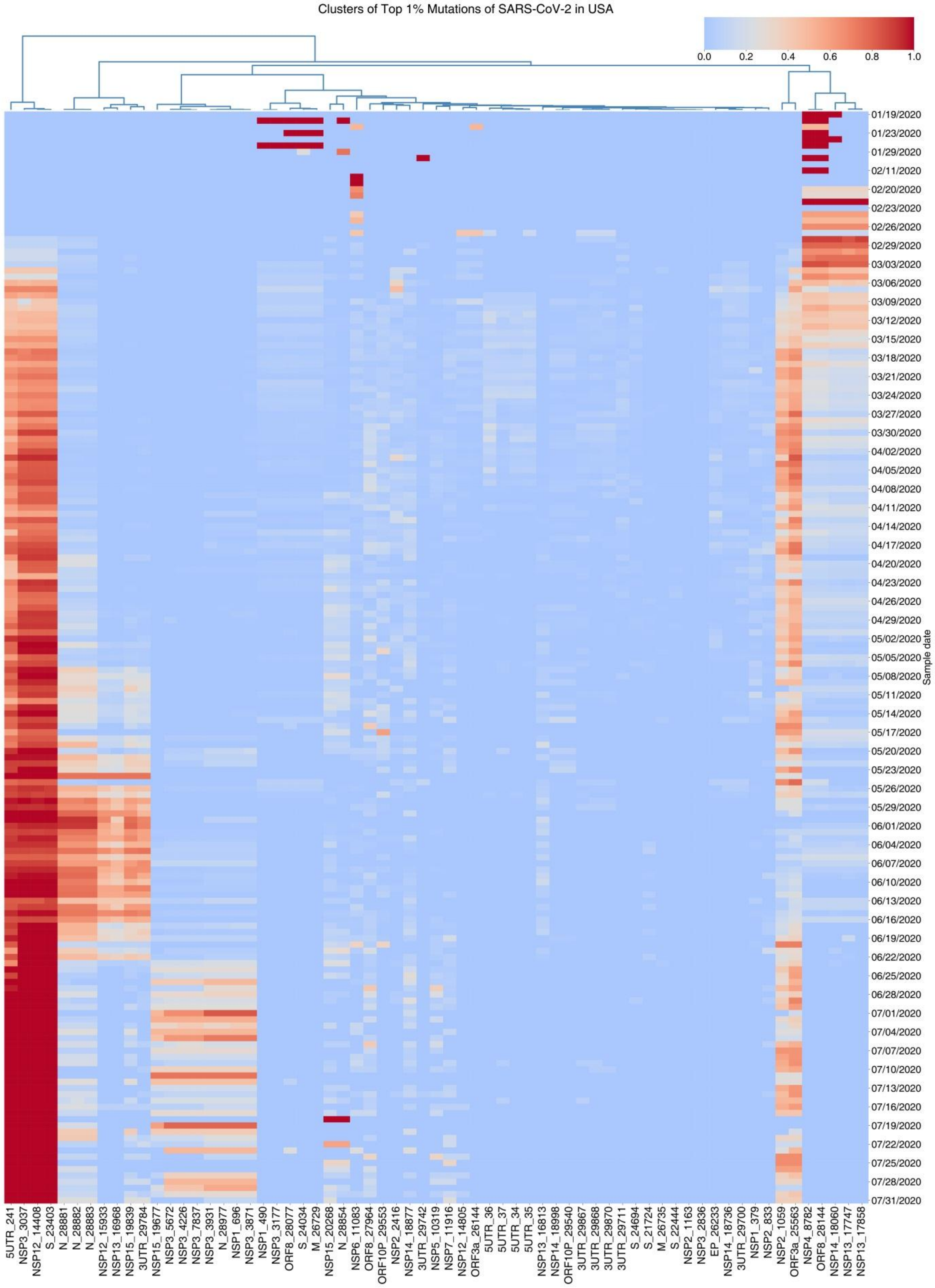


**Fig. S2 Ongoing mutations indicate potential co-mutations of SARS-CoV-2 from U.S. samples.** The number of sequences per collection date is shown in Table S6. Top 1% high-frequency mutations consist of 65 sites.There are also about 25 mutations clustered into several potential co-mutation patterns.

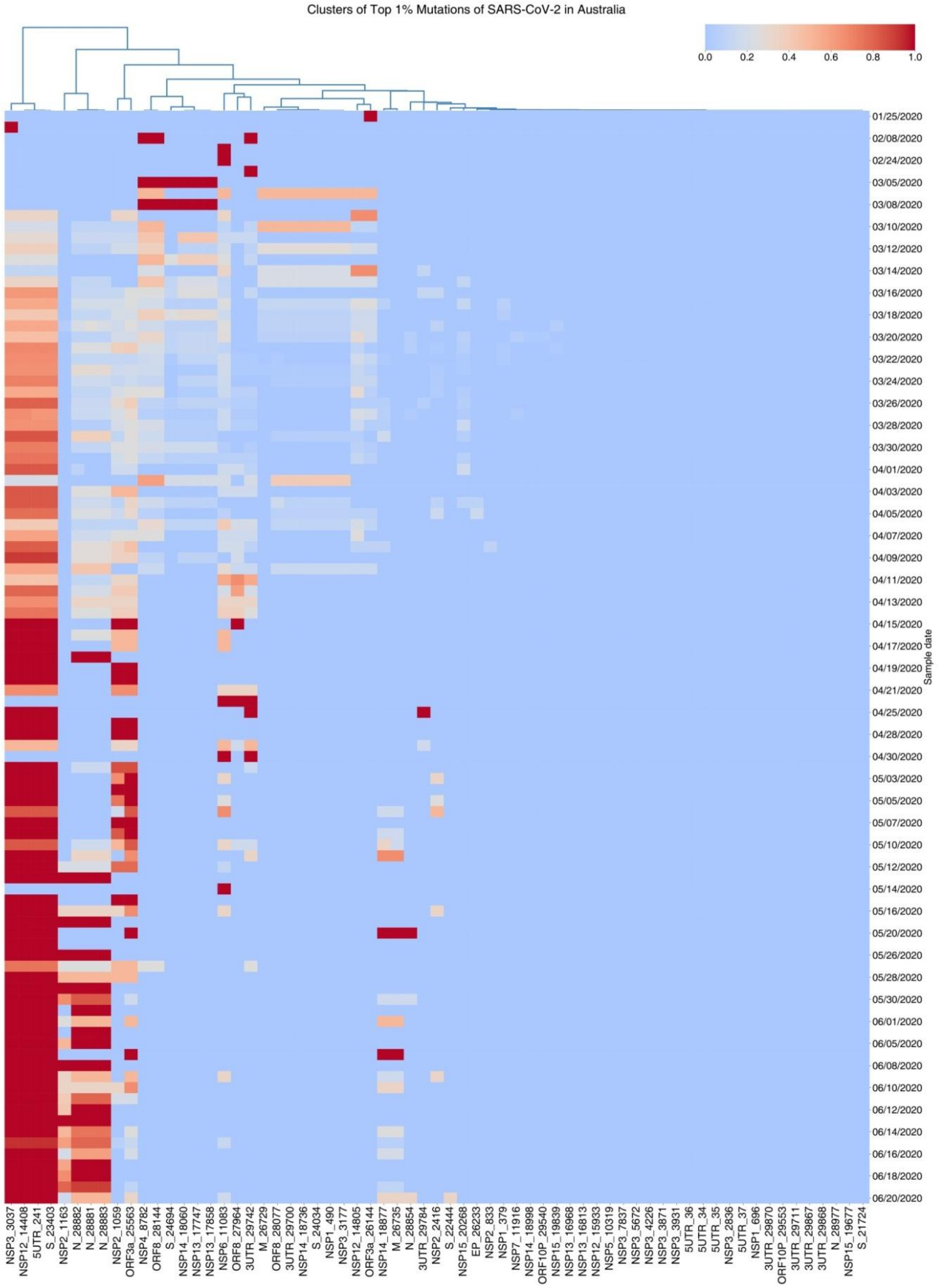


**Fig. S3 Ongoing mutations indicate potential co-mutations of SARS-CoV-2 from Australian samples.** The number of sequences per collection date is shown in Table S7. Top 1% high-frequency mutations consist of 65 sites. There are three possible co-mutation patterns consisting of the top 9 high-frequency mutations.

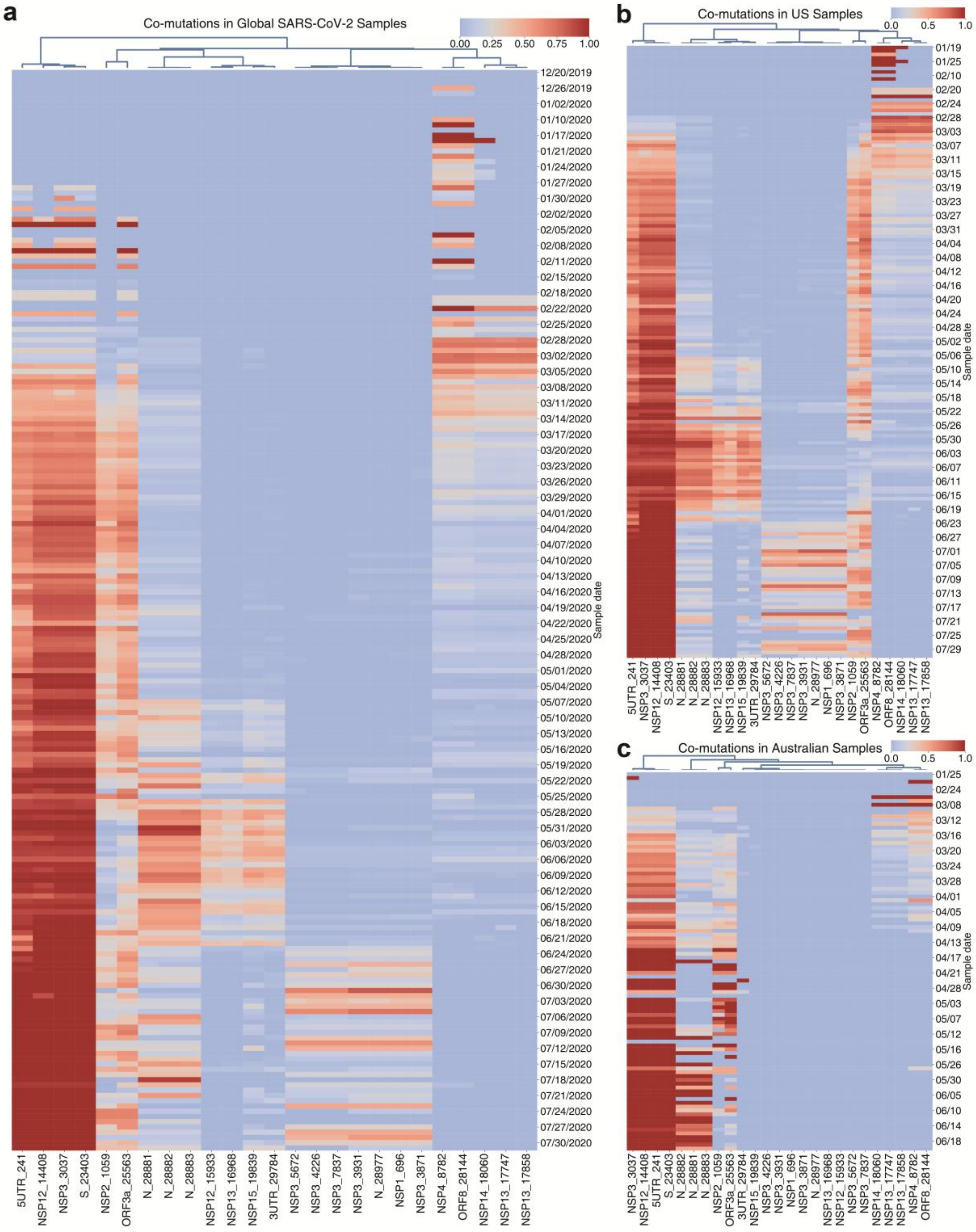


**Fig. S4 Ongoing mutations converge into potential co-mutation patterns in SARS-CoV-2 samples. a** Landscape of co-mutations of the global SARS-CoV-2 samples. The collection dates of these coronavirus strains range from Dec. 20, 2019 to Jul. 31, 2020. **b**,**c** Landscapes of co-mutations of SARS-CoV-2 samples in USA and in Australia respectively. These 25 sites are almost the top high-frequency mutation sites of SARS-CoV-2. The clusters of the mutations show that several potential co-mutation patterns with different evolutionary trends in not only global samples, but U.S. and Australian ones.

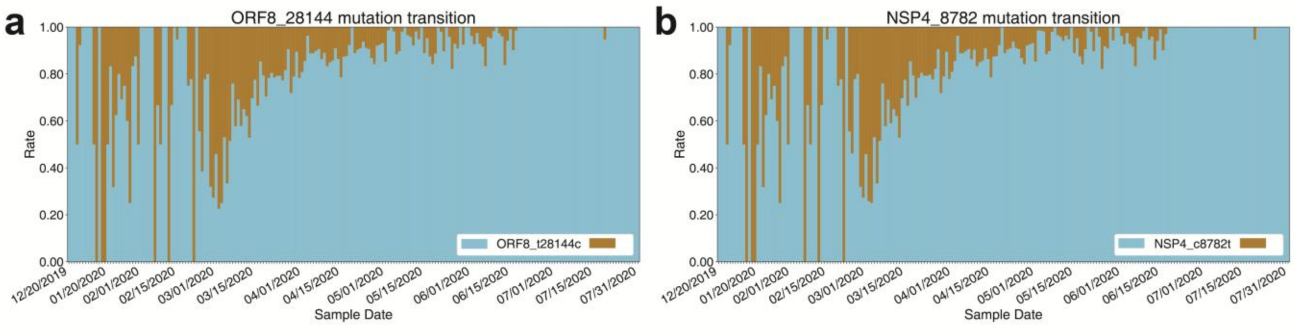


**Fig. S5 Mutation transitions of ORF3a_g25563t and NSP2_c1059t.** The mutational trends of these two mutations are similar.

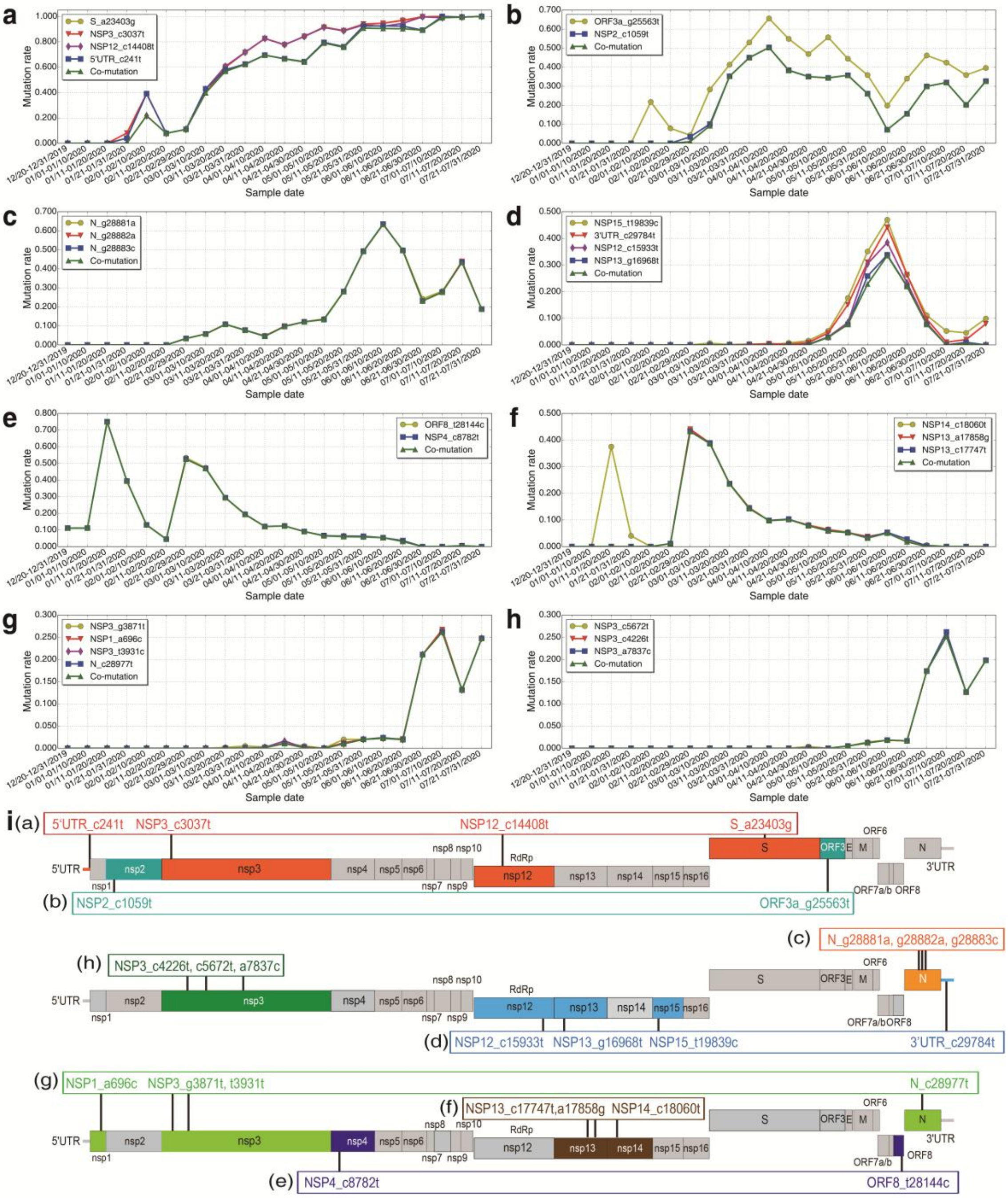


**Fig. S6 Evolutionary trajectories of potential co-mutations. a** Mutation and co-mutation trends of four sites. Mutations S_a23403g, NSP3_c3037t, NSP12_c14408t, and 5'UTR_c241t all became the dominant variants and such four sites almost co-occur (green curve) over time. **b** The co-mutation rate of ORF3a_g25563t and NSP2_c1059t is nearly the mutation rate of NSP2_c1059t in each time span. **c** The mutations and potential co-mutations of three successive sites almost share an identical trend curve. The co-mutation rates of both **e** and **f** decrease overall, while those of **g** and **h** totally increase. Differently, co-occurrence mutations in **d** appear and increase, then decrease and disappear. **i** The positions of eight potential co-mutation patterns including 25 sites in SARS-CoV-2 genome. Such patterns that reside in multi-genes may imply the potential interactions between/among sites and the evolutionary features.

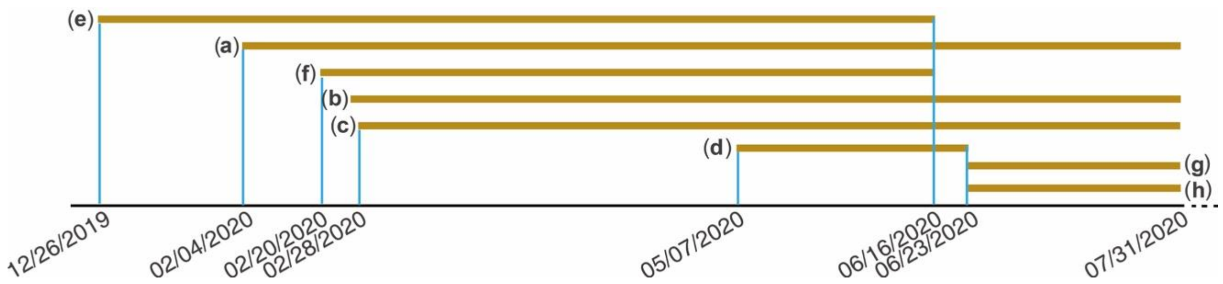


**Fig. S7 Highly dynamic co-mutation patterns.** The black line at the bottom of the figure is the time axis. Each golden line represents a potential co-mutation pattern that corresponds to Fig. S6.

**Table S1** Possible animal hosts of the direct progenitor of variant B.1.1.7. The strain labled by a star corresponds with the star variant as shown in main text Fig. 1a.

| Strain ID | Host | Collection Location | Collection Date | Mutation |
|---|---|---|---|---|
| EPI_ISL_752700 (*) | Homo | USA,Hawaii | 03/29/2020 | D614G,P681H |
| EPI_ISL_699508 | Dog | USA,Texas,Brazos County | 07/28/2020 | D614G,P681H |
| MT724346 | Tiger | USA | 04/04/2020 | D614G,T716I |
| EPI_ISL_641506 | Mink | Denmark | 08/13/2020 | 69-70del,D614G |

**Table S2** Comparative analysis of three possible animal hosts of the direct progenitor of variant B.1.1.7.

| Strain ID | Canidae (Dog) | Felidae (Tiger) | Mustelidae (Mink) |
|---|---|---|---|
| Mutation Edit Distance | 0 | 1 | 2 |
| Spike Similarity | 0.0011 | 0.0015 | 0.0012 |
| Collection Date | 07/28/2020 | 04/04/2020 | 08/13/2020 |
| Collection Location | USA | USA | Denmark |
| #Possible Transmission Origins (Homo) | 77 | 12 | 45 |

**Table S3** Quantitative analysis of three possible animal hosts based on Table S2. The smaller the value, the more similar with the star variant.

| Strain ID | Canidae (Dog) | Felidae (Tiger) | Mustelidae(Mink) |
|---|---|---|---|
| Mutation Edit Distance | 1 | 2 | 3 |
| Spike Similarity | 1 | 3 | 2 |
| Collection Date | 2 | 1 | 3 |
| Collection Location | 1 | 1 | 2 |
| #Possible Transmission Origins (Homo) | 1 | 3 | 2 |
| Sum | 6 | 10 | 12 |

**Table S4** Top 1% high-frequency mutations. There are 25 mutations formed eight potential co-mutation patterns. The mutations in the same background color indicate that they co-mutate with each other.

| Site | Ref. Site | Mutant | Mut. Rate | Ref. Codon | Mut. Codon | Ref. AA | Mut. AA |
|---|---|---|---|---|---|---|---|
| S_23403 | a | g | 0.7764 | gat | ggt | D | G |
| NSP3_3037 | c | t | 0.7744 | ttc | ttt | F | F |
| NSP12_14408 | c | t | 0.7720 | cta | tta | L | L |
| 5UTR_241 | c | t | 0.6964 | / | / | / | / |
| ORF3a_25563 | g | t | 0.4499 | cag | cat | Q | H |
| NSP2_1059 | c | t | 0.3303 | acc | atc | T | I |
| N_28881 | g | a | 0.1863 | agg | aag | R | K |
| N_28882 | g | a | 0.1856 | agg | aga | R | R |
| N_28883 | g | c | 0.1855 | gga | cga | G | R |
| ORF8_28144 | t | c | 0.1500 | tca | tta | L | S |
| NSP4_8782 | c | t | 0.1497 | agc | agt | S | S |
| NSP14_18060 | c | t | 0.1171 | ctc | ctt | L | L |
| NSP13_17858 | a | g | 0.1161 | tat | tgt | Y | C |
| NSP13_17747 | c | t | 0.1160 | cct | ctt | P | L |
| NSP15_19839 | t | c | 0.0793 | ata | aca | I | T |
| NSP14_18877 | c | t | 0.0778 | gtc | gtt | V | V |
| 3UTR_29784 | c | t | 0.0723 | / | / | / | / |
| NSP12_15933 | c | t | 0.0597 | acc | atc | T | I |
| ORF8_27964 | c | t | 0.0593 | gtc | gtt | V | V |
| NSP6_11083 | g | t | 0.0544 | ttg | ttt | L | F |
| NSP13_16968 | g | t | 0.0534 | agc | atc | S | I |
| N_28854 | c | t | 0.0501 | tca | tta | S | L |
| NSP15_20268 | a | g | 0.0454 | tag | tgg | terminator | G |
| ORF10P_29553 | g | a | 0.0416 | aag | aaa | K | K |
| NSP2_2416 | c | t | 0.0354 | tac | tat | Y | Y |
| 5UTR_36 | c | t | 0.0349 | / | / | / | / |
| M_26735 | c | t | 0.0283 | cag | tag | Q | terminator |
| 5UTR_35 | a | t | 0.0243 | / | / | / | / |
| 5UTR_34 | a | t | 0.0224 | / | / | / | / |
| NSP3_3871 | g | t | 0.0224 | aag | aat | K | N |
| NSP2_1163 | a | t | 0.0217 | att | ttt | I | F |
| NSP12_14805 | c | t | 0.0216 | act | att | T | I |
| NSP1_696 | a | c | 0.0205 | gac | gcc | D | G |
| NSP3_3931 | t | c | 0.0205 | gtt | gtc | V | V |
| N_28977 | c | t | 0.0204 | tct | ttt | S | F |
| NSP7_11916 | c | t | 0.0200 | tca | tta | S | L |
| S_22444 | c | t | 0.0197 | gac | gat | D | D |
| NSP1_379 | c | a | 0.0188 | gtc | gta | V | V |
| ORF8_28077 | g | c | 0.0185 | cgt | cct | R | P |
| 5UTR_37 | c | a | 0.0183 | / | / | / | / |

| | | | | | | | |
|---|---|---|---|---|---|---|---|
| ORF3a_26144 | g | t | 0.0183 | gtt | ttt | V | F |
| 3UTR_29700 | a | g | 0.0177 | / | / | / | / |
| NSP3_3177 | c | t | 0.0174 | cct | ctt | P | L |
| S_24034 | c | t | 0.0174 | aac | aat | N | N |
| NSP1_490 | t | a | 0.0169 | gat | gaa | D | E |
| NSP3_2836 | c | t | 0.0168 | tgc | tgt | C | C |
| NSP3_5672 | c | t | 0.0168 | cct | tct | P | S |
| NSP3_4226 | c | t | 0.0167 | cca | tca | P | S |
| NSP3_7837 | a | c | 0.0167 | tta | ttc | L | F |
| NSP14_18736 | t | c | 0.0163 | gat | gac | D | D |
| NSP2_833 | t | c | 0.0161 | ttc | ctc | F | L |
| M_26729 | t | c | 0.0161 | tgt | cgt | C | R |
| EP_26233 | g | t | 0.0150 | atg | att | M | I |
| 3UTR_29870 | c | a | 0.0141 | / | / | / | / |
| NSP13_16813 | g | a | 0.0133 | aag | aaa | K | K |
| NSP15_19677 | g | t | 0.0132 | agg | atg | R | M |
| NSP5_10319 | c | t | 0.0126 | ctt | ttt | L | F |
| 3UTR_29868 | g | a | 0.0122 | / | / | / | / |
| 3UTR_29711 | g | t | 0.0120 | / | / | / | / |
| 3UTR_29742 | g | t | 0.0119 | / | / | / | / |
| ORF10P_29540 | g | a | 0.0111 | tgc | tac | C | Y |
| NSP14_18998 | c | t | 0.0110 | cat | tat | H | Y |
| S_24694 | a | t | 0.0104 | gga | ggt | G | G |
| 3UTR_29867 | t | a | 0.0104 | / | / | / | / |
| S_21724 | g | t | 0.0100 | ttg | ttt | L | F |

**Table S5** Sample numbers of collected dates of SARS-CoV-2 strains. These samples were used for Fig. S1 and Fig. S4a.

| Collection date | Sample number |
|---|---|
| 12/20/2019 | 1 |
| 12/21/2019 | 1 |
| 12/23/2019 | 1 |
| 12/26/2019 | 2 |
| 12/30/2019 | 13 |
| 01/01/2020 | 2 |
| 01/02/2020 | 2 |
| 01/03/2020 | 1 |
| 01/08/2020 | 2 |
| 01/10/2020 | 2 |
| 01/11/2020 | 1 |
| 01/13/2020 | 1 |
| 01/17/2020 | 1 |
| 01/19/2020 | 3 |
| 01/20/2020 | 2 |
| 01/21/2020 | 6 |
| 01/22/2020 | 22 |
| 01/23/2020 | 8 |
| 01/24/2020 | 5 |
| 01/25/2020 | 13 |
| 01/26/2020 | 8 |
| 01/27/2020 | 5 |
| 01/28/2020 | 8 |
| 01/29/2020 | 12 |
| 01/30/2020 | 8 |
| 01/31/2020 | 4 |
| 02/01/2020 | 2 |
| 02/02/2020 | 2 |
| 02/03/2020 | 3 |
| 02/04/2020 | 1 |
| 02/05/2020 | 3 |
| 02/06/2020 | 1 |
| 02/07/2020 | 3 |
| 02/08/2020 | 2 |
| 02/09/2020 | 1 |
| 02/10/2020 | 5 |
| 02/11/2020 | 1 |
| 02/13/2020 | 3 |
| 02/14/2020 | 1 |
| 02/15/2020 | 19 |
| 02/16/2020 | 21 |
| 02/17/2020 | 32 |
| 02/18/2020 | 8 |
| 02/20/2020 | 4 |
| 02/21/2020 | 9 |
| 02/22/2020 | 3 |
| 02/23/2020 | 2 |
| 02/24/2020 | 9 |
| 02/25/2020 | 13 |
| 02/26/2020 | 9 |
| 02/27/2020 | 15 |
| 02/28/2020 | 25 |
| 02/29/2020 | 33 |
| 03/01/2020 | 24 |
| 03/02/2020 | 31 |
| 03/03/2020 | 24 |
| 03/04/2020 | 32 |
| 03/05/2020 | 54 |
| 03/06/2020 | 33 |
| 03/07/2020 | 54 |
| 03/08/2020 | 45 |
| 03/09/2020 | 55 |
| 03/10/2020 | 83 |
| 03/11/2020 | 80 |
| 03/12/2020 | 116 |
| 03/13/2020 | 229 |
| 03/14/2020 | 155 |
| 03/15/2020 | 147 |
| 03/16/2020 | 167 |
| 03/17/2020 | 157 |
| 03/18/2020 | 165 |
| 03/19/2020 | 213 |
| 03/20/2020 | 226 |
| 03/21/2020 | 174 |
| 03/22/2020 | 130 |
| 03/23/2020 | 207 |
| 03/24/2020 | 241 |
| 03/25/2020 | 242 |
| 03/26/2020 | 185 |
| 03/27/2020 | 202 |
| 03/28/2020 | 178 |
| 03/29/2020 | 95 |
| 03/30/2020 | 162 |
| 03/31/2020 | 194 |
| 04/01/2020 | 231 |
| 04/02/2020 | 214 |
| 04/03/2020 | 191 |
| 04/04/2020 | 133 |
| 04/05/2020 | 99 |
| 04/06/2020 | 167 |
| 04/07/2020 | 128 |

| | |
|---|---|
| 04/08/2020 | 124 |
| 04/09/2020 | 73 |
| 04/10/2020 | 66 |
| 04/11/2020 | 60 |
| 04/12/2020 | 49 |
| 04/13/2020 | 111 |
| 04/14/2020 | 88 |
| 04/15/2020 | 70 |
| 04/16/2020 | 87 |
| 04/17/2020 | 73 |
| 04/18/2020 | 52 |
| 04/19/2020 | 25 |
| 04/20/2020 | 54 |
| 04/21/2020 | 83 |
| 04/22/2020 | 122 |
| 04/23/2020 | 97 |
| 04/24/2020 | 79 |
| 04/25/2020 | 32 |
| 04/26/2020 | 53 |
| 04/27/2020 | 108 |
| 04/28/2020 | 114 |
| 04/29/2020 | 148 |
| 04/30/2020 | 160 |
| 05/01/2020 | 122 |
| 05/02/2020 | 70 |
| 05/03/2020 | 66 |
| 05/04/2020 | 110 |
| 05/05/2020 | 95 |
| 05/06/2020 | 78 |
| 05/07/2020 | 48 |
| 05/08/2020 | 24 |
| 05/09/2020 | 29 |
| 05/10/2020 | 24 |
| 05/11/2020 | 51 |
| 05/12/2020 | 54 |
| 05/13/2020 | 58 |
| 05/14/2020 | 57 |
| 05/15/2020 | 36 |
| 05/16/2020 | 21 |
| 05/17/2020 | 8 |
| 05/18/2020 | 38 |
| 05/19/2020 | 53 |
| 05/20/2020 | 27 |
| 05/21/2020 | 48 |
| 05/22/2020 | 29 |
| 05/23/2020 | 24 |
| 05/24/2020 | 24 |
| 05/25/2020 | 28 |
| 05/26/2020 | 98 |
| 05/27/2020 | 110 |
| 05/28/2020 | 37 |
| 05/29/2020 | 53 |
| 05/30/2020 | 26 |
| 05/31/2020 | 22 |
| 06/01/2020 | 79 |
| 06/02/2020 | 123 |
| 06/03/2020 | 80 |
| 06/04/2020 | 42 |
| 06/05/2020 | 96 |
| 06/06/2020 | 24 |
| 06/07/2020 | 47 |
| 06/08/2020 | 124 |
| 06/09/2020 | 59 |
| 06/10/2020 | 39 |
| 06/11/2020 | 74 |
| 06/12/2020 | 74 |
| 06/13/2020 | 43 |
| 06/14/2020 | 34 |
| 06/15/2020 | 110 |
| 06/16/2020 | 81 |
| 06/17/2020 | 67 |
| 06/18/2020 | 114 |
| 06/19/2020 | 61 |
| 06/20/2020 | 63 |
| 06/21/2020 | 23 |
| 06/22/2020 | 61 |
| 06/23/2020 | 47 |
| 06/24/2020 | 42 |
| 06/25/2020 | 31 |
| 06/26/2020 | 40 |
| 06/27/2020 | 27 |
| 06/28/2020 | 27 |
| 06/29/2020 | 30 |
| 06/30/2020 | 23 |
| 07/01/2020 | 6 |
| 07/02/2020 | 15 |
| 07/03/2020 | 16 |
| 07/04/2020 | 9 |
| 07/05/2020 | 11 |
| 07/06/2020 | 39 |
| 07/07/2020 | 29 |
| 07/08/2020 | 17 |
| 07/09/2020 | 23 |
| 07/10/2020 | 26 |
| 07/11/2020 | 5 |
| 07/12/2020 | 11 |
| 07/13/2020 | 12 |
| 07/14/2020 | 32 |

| | |
|---|---|
| 07/15/2020 | 66 |
| 07/16/2020 | 15 |
| 07/17/2020 | 10 |
| 07/18/2020 | 14 |
| 07/19/2020 | 19 |
| 07/20/2020 | 14 |
| 07/21/2020 | 16 |
| 07/22/2020 | 12 |
| 07/23/2020 | 4 |
| 07/24/2020 | 9 |
| 07/25/2020 | 3 |
| 07/26/2020 | 5 |
| 07/27/2020 | 15 |
| 07/28/2020 | 6 |
| 07/29/2020 | 9 |
| 07/30/2020 | 14 |
| 07/31/2020 | 8 |

**Table S6** Sample numbers of collected dates of SARS-CoV-2 strains in USA. These samples were used for Fig. S2 and Fig. S4b.

| Collection date | Sample number |
|---|---|
| 01/19/2020 | 3 |
| 01/21/2020 | 1 |
| 01/22/2020 | 2 |
| 01/23/2020 | 1 |
| 01/25/2020 | 2 |
| 01/28/2020 | 1 |
| 01/29/2020 | 4 |
| 02/06/2020 | 1 |
| 02/10/2020 | 1 |
| 02/11/2020 | 1 |
| 02/17/2020 | 8 |
| 02/18/2020 | 6 |
| 02/20/2020 | 3 |
| 02/21/2020 | 7 |
| 02/22/2020 | 2 |
| 02/23/2020 | 1 |
| 02/24/2020 | 5 |
| 02/25/2020 | 2 |
| 02/26/2020 | 3 |
| 02/27/2020 | 7 |
| 02/28/2020 | 19 |
| 02/29/2020 | 30 |
| 03/01/2020 | 18 |
| 03/02/2020 | 30 |
| 03/03/2020 | 21 |
| 03/04/2020 | 26 |
| 03/05/2020 | 50 |
| 03/06/2020 | 33 |
| 03/07/2020 | 48 |
| 03/08/2020 | 39 |
| 03/09/2020 | 36 |
| 03/10/2020 | 52 |
| 03/11/2020 | 51 |
| 03/12/2020 | 92 |
| 03/13/2020 | 208 |
| 03/14/2020 | 128 |
| 03/15/2020 | 108 |
| 03/16/2020 | 132 |
| 03/17/2020 | 119 |
| 03/18/2020 | 109 |
| 03/19/2020 | 174 |
| 03/20/2020 | 158 |
| 03/21/2020 | 128 |
| 03/22/2020 | 81 |
| 03/23/2020 | 145 |
| 03/24/2020 | 194 |
| 03/25/2020 | 206 |
| 03/26/2020 | 156 |
| 03/27/2020 | 153 |
| 03/28/2020 | 150 |
| 03/29/2020 | 65 |
| 03/30/2020 | 126 |
| 03/31/2020 | 167 |
| 04/01/2020 | 208 |
| 04/02/2020 | 202 |
| 04/03/2020 | 170 |
| 04/04/2020 | 120 |
| 04/05/2020 | 84 |
| 04/06/2020 | 154 |
| 04/07/2020 | 111 |
| 04/08/2020 | 107 |
| 04/09/2020 | 61 |
| 04/10/2020 | 55 |
| 04/11/2020 | 40 |
| 04/12/2020 | 38 |
| 04/13/2020 | 100 |
| 04/14/2020 | 77 |
| 04/15/2020 | 67 |
| 04/16/2020 | 81 |
| 04/17/2020 | 69 |
| 04/18/2020 | 47 |
| 04/19/2020 | 22 |
| 04/20/2020 | 51 |
| 04/21/2020 | 77 |
| 04/22/2020 | 110 |
| 04/23/2020 | 94 |
| 04/24/2020 | 75 |
| 04/25/2020 | 27 |
| 04/26/2020 | 38 |
| 04/27/2020 | 93 |
| 04/28/2020 | 90 |
| 04/29/2020 | 111 |
| 04/30/2020 | 153 |
| 05/01/2020 | 116 |
| 05/02/2020 | 27 |
| 05/03/2020 | 43 |
| 05/04/2020 | 103 |
| 05/05/2020 | 72 |
| 05/06/2020 | 66 |
| 05/07/2020 | 43 |
| 05/08/2020 | 23 |

| | |
|---|---|
| 05/09/2020 | 19 |
| 05/10/2020 | 8 |
| 05/11/2020 | 44 |
| 05/12/2020 | 45 |
| 05/13/2020 | 55 |
| 05/14/2020 | 52 |
| 05/15/2020 | 32 |
| 05/16/2020 | 15 |
| 05/17/2020 | 5 |
| 05/18/2020 | 36 |
| 05/19/2020 | 49 |
| 05/20/2020 | 25 |
| 05/21/2020 | 44 |
| 05/22/2020 | 29 |
| 05/23/2020 | 9 |
| 05/24/2020 | 4 |
| 05/25/2020 | 27 |
| 05/26/2020 | 92 |
| 05/27/2020 | 81 |
| 05/28/2020 | 33 |
| 05/29/2020 | 52 |
| 05/30/2020 | 14 |
| 05/31/2020 | 13 |
| 06/01/2020 | 57 |
| 06/02/2020 | 72 |
| 06/03/2020 | 54 |
| 06/04/2020 | 41 |
| 06/05/2020 | 67 |
| 06/06/2020 | 21 |
| 06/07/2020 | 18 |
| 06/08/2020 | 94 |
| 06/09/2020 | 53 |
| 06/10/2020 | 36 |
| 06/11/2020 | 40 |
| 06/12/2020 | 25 |
| 06/13/2020 | 17 |
| 06/14/2020 | 18 |
| 06/15/2020 | 61 |
| 06/16/2020 | 60 |
| 06/17/2020 | 41 |
| 06/18/2020 | 58 |
| 06/19/2020 | 47 |
| 06/20/2020 | 46 |
| 06/21/2020 | 23 |
| 06/22/2020 | 59 |
| 06/23/2020 | 42 |
| 06/24/2020 | 36 |
| 06/25/2020 | 25 |
| 06/26/2020 | 37 |
| 06/27/2020 | 24 |
| 06/28/2020 | 26 |
| 06/29/2020 | 23 |
| 06/30/2020 | 23 |
| 07/01/2020 | 6 |
| 07/02/2020 | 13 |
| 07/03/2020 | 15 |
| 07/04/2020 | 8 |
| 07/05/2020 | 11 |
| 07/06/2020 | 15 |
| 07/07/2020 | 14 |
| 07/08/2020 | 17 |
| 07/09/2020 | 20 |
| 07/10/2020 | 25 |
| 07/11/2020 | 4 |
| 07/12/2020 | 11 |
| 07/13/2020 | 11 |
| 07/14/2020 | 25 |
| 07/15/2020 | 14 |
| 07/16/2020 | 13 |
| 07/17/2020 | 7 |
| 07/18/2020 | 1 |
| 07/19/2020 | 5 |
| 07/20/2020 | 5 |
| 07/21/2020 | 12 |
| 07/22/2020 | 12 |
| 07/23/2020 | 4 |
| 07/24/2020 | 9 |
| 07/25/2020 | 3 |
| 07/26/2020 | 5 |
| 07/27/2020 | 11 |
| 07/28/2020 | 6 |
| 07/29/2020 | 9 |
| 07/30/2020 | 14 |
| 07/31/2020 | 8 |

**Table S7** Sample numbers of collected dates of SARS-CoV-2 strains in Australia. These samples were used for Fig. S3 and Fig. S4c.

| Collection date | Sample number |
|---|---|
| 01/25/2020 | 1 |
| 01/30/2020 | 4 |
| 02/08/2020 | 1 |
| 02/21/2020 | 2 |
| 02/24/2020 | 1 |
| 03/03/2020 | 1 |
| 03/05/2020 | 1 |
| 03/07/2020 | 2 |
| 03/08/2020 | 1 |
| 03/09/2020 | 3 |
| 03/10/2020 | 10 |
| 03/11/2020 | 7 |
| 03/12/2020 | 11 |
| 03/13/2020 | 8 |
| 03/14/2020 | 9 |
| 03/15/2020 | 9 |
| 03/16/2020 | 8 |
| 03/17/2020 | 16 |
| 03/18/2020 | 24 |
| 03/19/2020 | 23 |
| 03/20/2020 | 41 |
| 03/21/2020 | 38 |
| 03/22/2020 | 39 |
| 03/23/2020 | 38 |
| 03/24/2020 | 31 |
| 03/25/2020 | 27 |
| 03/26/2020 | 21 |
| 03/27/2020 | 33 |
| 03/28/2020 | 20 |
| 03/29/2020 | 19 |
| 03/30/2020 | 25 |
| 03/31/2020 | 16 |
| 04/01/2020 | 12 |
| 04/02/2020 | 5 |
| 04/03/2020 | 12 |
| 04/04/2020 | 12 |
| 04/05/2020 | 13 |
| 04/06/2020 | 10 |
| 04/07/2020 | 12 |
| 04/08/2020 | 11 |
| 04/09/2020 | 11 |
| 04/10/2020 | 7 |
| 04/11/2020 | 9 |
| 04/12/2020 | 5 |
| 04/13/2020 | 3 |
| 04/14/2020 | 8 |
| 04/15/2020 | 1 |
| 04/16/2020 | 4 |
| 04/17/2020 | 2 |
| 04/18/2020 | 1 |
| 04/19/2020 | 1 |
| 04/20/2020 | 1 |
| 04/21/2020 | 3 |
| 04/22/2020 | 1 |
| 04/25/2020 | 1 |
| 04/27/2020 | 2 |
| 04/28/2020 | 1 |
| 04/29/2020 | 6 |
| 04/30/2020 | 1 |
| 05/02/2020 | 7 |
| 05/03/2020 | 3 |
| 05/04/2020 | 3 |
| 05/05/2020 | 4 |
| 05/06/2020 | 6 |
| 05/07/2020 | 2 |
| 05/09/2020 | 6 |
| 05/10/2020 | 6 |
| 05/11/2020 | 3 |
| 05/12/2020 | 9 |
| 05/13/2020 | 1 |
| 05/14/2020 | 3 |
| 05/15/2020 | 2 |
| 05/16/2020 | 3 |
| 05/19/2020 | 4 |
| 05/20/2020 | 1 |
| 05/23/2020 | 1 |
| 05/26/2020 | 1 |
| 05/27/2020 | 4 |
| 05/28/2020 | 4 |
| 05/29/2020 | 1 |
| 05/30/2020 | 6 |
| 05/31/2020 | 1 |
| 06/01/2020 | 4 |
| 06/03/2020 | 1 |
| 06/05/2020 | 2 |
| 06/06/2020 | 2 |
| 06/08/2020 | 1 |
| 06/09/2020 | 6 |
| 06/10/2020 | 3 |
| 06/11/2020 | 5 |
| 06/12/2020 | 7 |

| | | | | |
|---|---|---|---|---|
| 06/13/2020 | 5 | | 06/17/2020 | 7 |
| 06/14/2020 | 4 | | 06/18/2020 | 6 |
| 06/15/2020 | 15 | | 06/19/2020 | 10 |
| 06/16/2020 | 5 | | 06/20/2020 | 6 |
